# Changes in Cellular Crosstalk between Skeletal Muscle Myoblasts and Bone Osteoblasts with Aging

**DOI:** 10.1101/2021.04.26.441458

**Authors:** Jonathan A. Doering, Carly E. Britt, Gregory S. Sawicki, Jacqueline H. Cole

## Abstract

Musculoskeletal function declines with aging, resulting in an increased incidence of trips and falls. Both bone and muscle experience age-related losses in tissue mass that alter their mechanical interactions in a well characterized manner, but changes in the biochemical interactions between bone and muscle with aging are not well understood. Of note, insulin-like growth factor 1 (IGF-1), a potent growth factor for bone and muscle, can be negatively altered with aging and may help explain losses in these tissues. We recently developed a co-culture system for simultaneous growth of bone mesenchymal stem cells (MSCs) and muscle satellite cells (SCs) to investigate the biochemical crosstalk between the two cell types. Here, we utilized an aging rat model to study cellular changes between young and old rat MSCs and SCs, in particular whether 1) young MSCs and SCs have increased proliferation and differentiation compared to old MSCs and SCs; 2) young cells have increased IGF-1 and collagen expression as a measure of crosstalk compared to old cells; and 3) young cells can mitigate the aging phenotype of old cells in co-culture. Rat MSCs and SCs were either mono- or co-cultured in Transwell^®^ plates, grown to confluence, and allowed to differentiate for 14 days. Across the 14 days, cell proliferation was measured, with differentiation and crosstalk measurements evaluated at 14 days. The results suggest that in both young and old, proliferation is greater in mono-cultures compared to co-cultures, yet age and cell type did not have a significant effect. Differentiation did not differ between young and old cells, yet MSCs and SCs demonstrated the greatest amount of differentiation in co-culture. Finally, age, cell type, and culture type did not have a significant effect on collagen or IGF-1 expression. These results suggest co-culture may have a controlling effect, with the two cell types acting together to promote differentiation more than in mono-cultures, yet this response was not altered by age. In general, results for old cells had higher variability, suggesting a wider variety in the aging phenotypes demonstrated in these animals. This study was the first to use this rat aging model to investigate changes between bone and skeletal muscle cells, however further investigations are required to determine what signaling changes occur in response to age. Determining these signaling changes could lead to new targets for mitigating the progression of aging.

## Introduction

Musculoskeletal performance declines with aging, resulting in an increased incidence of trips and falls in the elderly.^1^ These changes are compounded by concomitant losses in bone mass (osteopenia, osteoporosis) and skeletal muscle mass (sarcopenia), leading to high rates of fracture and muscle injury in the elderly population.^3,4^ Under the mechanostat paradigm, these detrimental muscle changes, which manifest in decreased muscle loading on the skeleton, will induce negative bone adaptation and bone loss.^2^ Aged individuals with decreased bone mass (osteopenia) often also present with reduced muscle mass and function (sarcopenia).^3,4^ However, although elderly populations often experience both osteopenia and sarcopenia, age-related declines in either bone or muscle mass cannot fully explain the extent of functional declines in the other tissue, supporting the idea that bone-muscle interactions are not solely mechanical in nature. Indeed, the emergent thought is that biochemical interactions also play an important role in bone and muscle tissue health and function, in particular the direct communication between the two through paracrine signaling pathways, or cell-cell *crosstalk*.^5,6^ Improving our understanding of how bone-muscle crosstalk changes with aging could inform the development of targeted therapies to mitigate age-related losses in musculoskeletal function.

Age-related changes in biochemical signaling are well characterized for bone and muscle individually. In muscle, satellite cells (SCs) responsible for muscle repair have decreased activity from overuse, and signaling molecules such as inflammatory and growth factors are reduced.^7,8^ Similarly, bone loss associated with aging is the result of reduced functional ability of bone osteoblasts to produce new bone, as well as a loss of signaling factors.^9^

While these signaling changes have been investigated in either muscle or bone alone, the investigation of the signaling changes in tandem have been under characterized. The concurrent age-related changes in bone and muscle signaling are not fully understood, although the importance of this crosstalk has been established in several key studies. In elegant parabiosis studies, in which the vascular systems for young and old rat pairs were surgically merged, old rats experienced some muscle rejuvenation, including increased muscle mass and improved muscle fiber structure, although specific factors from the young rats stimulating this change remained unidentified.^17,18^ Further, *in vivo* mouse studies demonstrated faster healing of bone fractures in the presence of additional xenografted muscle tissue,^19,20^ but again the specific crosstalk factors stimulating tissue repair were ill-defined, where specific signaling factors were not discussed. Therefore, a more specific method is required to investigate the specific crosstalk changes. Our previous work optimized a co-culture system between SCs and bone mesenchymal stem cells (MSCs) to investigate the optimal growth conditions for each cell type. Utilization of this system could lead to better understanding of the crosstalk between muscle and bone.

One signaling target for investigation in both muscle and bone is insulin-like growth factor 1 (IGF-1). IGF-1 is a potent growth factor secreted by the liver in response to the secretion of growth hormone, resulting in increased growth of tissues like bone and muscle.^10^ However, IGF-1 has also been demonstrated to be released by both muscle and bone to be used in an autocrine or paracrine fashion.^10,11^ Further, reduction in IGF-1 lead to reductions in both muscle and bone mass,^11-13^ and decreased the amount of collagen in these tissues.^14-16^ However, how IGF-1 is altered in the crosstalk between muscle and bone has not been investigated.

The purpose of this study was to characterize concurrent changes in bone and muscle cells and biochemical interactions with aging. Using our previously optimized co-culture system, we paired bone mesenchymal stem cells (MSCs) and muscle SCs from both young and old rats and examined cell viability, differentiation, and crosstalk using young-young, old-old, young-old, and old-young pairs. We hypothesized that in old cells, there would be decreased cell viability, differentiation, and crosstalk compared to young cells. Furthermore, we wanted to test the hypothesis that young factors could improve the aging phenotype as demonstrated by Conboy et al.^18^ By co-culturing young and old cells together we hypothesized that the young SCs or MSCs when cultured with the opposite old cell would be unchanged when compared to their regular young co-cultured with young cells, however, old cells under the influence of young cells would have increased cell viability, cell differentiation, and cell crosstalk when compared to old cells co-cultured with other old cells.

## Methods

### Animals and cell isolation

The Institutional Animal Care and Use Committee at North Carolina State University approved all animal procedures. F344 x Brown Norway F1 hybrid (F344BN) rats were used in this study as a model for aging, because they are commonly used in aging muscle function studies. Ten young (5-9 months old) and ten old (33-34 months old) F344BN rats (National Institutes of Aging, Bethesda, Maryland) were fed a normal diet and kept under a 12-hr light/12-hr dark cycle with free access to chow and water until euthanasia. After acclimation to the animal facility for one week, rats were euthanized by CO_2_ inhalation, followed by immediate dissection to remove the gastrocnemius and tibialis anterior (TA) muscles and the tibia and femur from one leg. Rat bone MSCs were isolated from the tibia and femur using a method modified from Zhang and Chan.^21^ Briefly, the bones were placed in warm MSC-specific growth medium (MSCGM) consisting of alpha-MEM (Gibco, Gaithersburg, MD), 10% fetal bovine serum (Gemini Bio-Products, West Sacramento, CA), and 100 U/mL penicillin-100 µg/mL streptomycin (Gibco). The bone ends were removed, and the remaining segments were flushed with MSCGM to extract the marrow and isolate the MSCs. The MSCs were plated on a single 120-mm plastic plate (Corning, Tewksbury, MA) coated with 2% gelatin (Fisher Scientific, Hampton, NH) and cultured with MSCGM in a humidified atmosphere at 37°C with 5% CO_2_. After 48 h, the nonadherent cells were washed out, and the cultures were expanded to passage 2.

SCs were extracted from the gastrocnemius and TA using a protocol from Danoviz and Yablonk-Reuveni.^22^ Briefly, the muscles were placed in warm SC-specific growth medium (SCGM) consisting of high glucose Dulbecco’s MEM (Gibco), 20% fetal bovine serum (Gemini Bioproducts), 10% horse serum (Gibco), and 100 U/mL penicillin-100 µg/mL streptomycin (Gibco). Tendon, fat, blood vessels, and connective tissue were removed, and the remaining muscle was cut into small (∼3 mm^3^) cubes. Muscle cubes were collected and added to a 1% Pronase solution (Pronase Protease, Millipore-Sigma, St. Louis, MO) in SCGM for one hour for digestion. The solution and remaining muscle were then collected and suspended in 10 mL of additional SCGM. The entire solution was subjected to mechanical trituration by passing the solution through a wide-bore pipette to liberate SCs. Finally, the triturated solution was passed through a cell strainer with a 70-µm pore size (Fisher Scientific), and SCs were collected and plated on a single 2% gelatin-coated 120-mm plate with SCGM in a humidified atmosphere at 37°C with 5% CO_2_. After 48 h, the nonadherent cells were washed out, and the cultures were expanded to passage 2.

### Co-culture design

After passage 2, cells were plated in both mono- and co-culture (Fig. 1). In mono-culture experiments, SCs and MSCs were each cultured at 20 x 10^3^ cells per well in a 12-well gelatin-coated plate, with each condition run in triplicate per animal. In co-culture experiments, SCs and MSCs were each cultured at 20 x 10^3^ cells per well in a 12-well Transwell^®^ gelatin-coated plate (Corning, Corning, NY), with SCs in the bottom well and MSCs in the insert, separated by the 0.4-µm pore membrane. Plates were allowed to grow to 50% confluence in SCGM for SCs and in MSCGM for MSCs. At 50% confluence, media was changed to myogenic media (MM) for SCs (containing high glucose Dulbecco’s MEM (Gibco), 1% horse serum (Gibco), and 100 U/mL penicillin-100 µg/mL streptomycin (Gibco)) and osteogenic media (OM) for MSCs (containing alpha-MEM (Gibco), 10% fetal bovine serum (Gibco), 50 mM ascorbic acid (Millipore-Sigma), 100 µM dexamethasone (Millipore-Sigma), 1 M β-glycerophosphate (Millipore-Sigma) and 100 U/mL penicillin-100 µg/mL streptomycin (Gibco)) to allow differentiation to occur. Differentiation continued for 14 days, at which point outcome metrics (below) were assessed.

**Figure 1.**
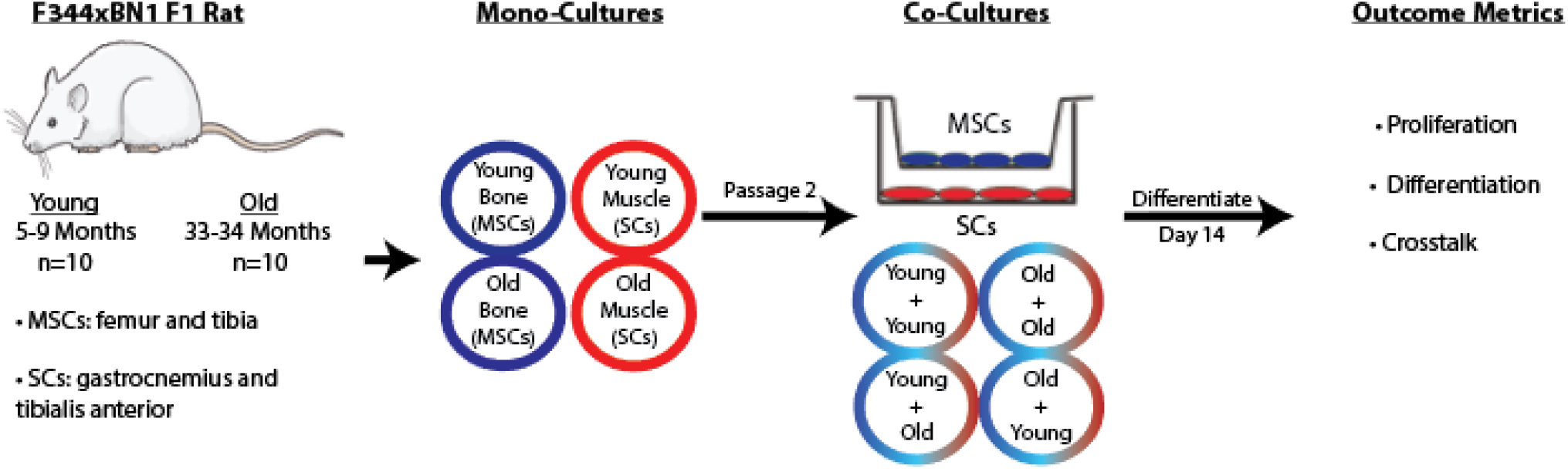
Study design. SCs and MSCs were isolated from rat tissue and cultivated in mono-culture and co-culture and allowed to differentiate for 14 days before outcome metrics were assessed.

### Cell proliferation

On differentiation days 0,3, 7, 10, and 14, an alamarBlue^®^ solution (ThermoFisher, Waltham, MA) was mixed into each well (90% media, 10% alamarBlue^®^), and the cells were incubated at 37°C. After 3 h, the solution was collected, and absorbance readings were taken at 570 and 600 nm (Synergy™ H1M, BioTek Instruments, Inc., Winooski, VT). For the co-cultures, the bottom well and insert of the Transwell^®^ plates were analyzed separately. The percent reduction in alamarBlue^®^ level, a measure of proliferative ability, was determined from the absorbance reading according to the manufacturer’s instructions.

### Cell differentiation

On differentiation day 14, cells were rinsed with phosphate buffered saline (PBS), fixed in 10% zinc-buffered formalin (VWR, Randor, PA) for 30 min, and then rinsed again with PBS. To assess mineral deposition for MSC differentiation, a 2% Alizarin Red S (Fisher) solution was applied to each sample for 5 min, rinsed with deionized water, and imaged using an iPhone 6S camera (Apple, Cupertino, CA) and EVOS^®^ XL light microscope (Life Technologies, Carlsbad, CA). Mineral deposition was quantified by solubilizing the Alizarin Red S stain with a 0.5 N HCl + 5% sodium dodecyl sulfate (SDS) solution for 30 min and then taking absorbance readings at 405 nm.

To assess SC differentiation, immunocytochemistry (ICC) procedures for skeletal myosin expression were performed. On differentiation day 14, cells were rinsed with DMEM, fixed in 10% zinc-buffered formalin for 10 min, and then rinsed with tris-buffered saline (TBS). A blocking solution of TBS with 2% normal goat serum (NGS) was applied to the cells and allowed to incubate at 4°C overnight. Cells were then allowed to warm to room temperature for 10 min. A rat-specific primary antibody for skeletal muscle myosin heavy chain (MA1-35718, ThermoFisher) was diluted 1:5 in TBS-NGS. Wells were rinsed with a TBS-0.05% Tween 20 (TW20, ThermoFisher) solution, and primary antibodies were applied. Cells were incubated for 1 h at room temperature, followed by an overnight incubation at 4°C in a humidified chamber with gentle shaking. Cells were again allowed to return to room temperature while the secondary antibody (AlexaFluor 584, A-21044, ThermoFisher) was diluted 1:10 in TBS-NGS. Cells were rinsed in TBS-TW20, and the secondary antibody was applied for 2 h at room temperature. Wells were aspirated and rinsed with TBS-TW20, and a diluted DAPI (ThermoFisher) solution was prepared (1 ug/mL diluted in TGS-NGS) and applied to the wells for 30 min at room temperature. Wells were rinsed with TBS-TW20 a final time and mounted with a drop of Vectashield (Vector Laboratories, Burlingame, CA) and a 25% glycerol solution in TBS. Cells were imaged (EVOS^®^ FL Auto, Life Technologies, Carlsbad, CA), and area and integrated density of the fluorescent regions were measured. SC and MSC differentiation were also assessed qualitatively based on cell morphology, looking for multi-nucleated and elongated cells with SCs and a fibroblastic appearance and mineral nodules with MSCs.

### Cellular crosstalk

Since IGF-1 aids growth in both bone and muscle, its expression was assessed in the conditioned media of both mono- and co-cultures at differentiation day 14 using an ELISA kit (ERIGF1, ThermoFisher), according to the manufacturer’s instructions. A standard curve was produced, and samples expressing IGF-1 levels outside of the standards’ range were omitted.

To assess the effect of aging on collagen production, ICC was performed to evaluate the amount of collagen I deposited by the cells. The ICC protocol was the same as for the cellular differentiation analysis, except with a rat-specific collagen I primary antibody (PA1-36145, ThermoFisher) and a secondary antibody (A-11034, ThermoFisher) used at a 1:10 dilution in TBS-NGS.

### Statistical analyses

Statistical analyses were performed with Prism (version 6.07; GraphPad Software, La Jolla, CA) using a significance level of 0.05. All results were averaged across the groups and expressed in the form of mean ± standard error of the mean due to unbalanced samples. The effects of age (young, old) and cell type (Mono-SC, Mono-MSC, Co-SC, Co-MSC) were examined using two-way ANOVAs. In cases where differentiation day was considered, a two-way ANOVA was performed with repeated measures. Post-hoc pairwise comparisons within each factor were evaluated with Tukey tests if interactions were previously found to be significant.

## Results

### Cell viability

Percent reduction in alamarBlue^®^ differed significantly by cell culture type at differentiation days 7, 10, and 14 (p < 0.05, Fig. 2), with mono-cultures demonstrating greater proliferation than co-cultures at these later time points, and with co-cultures demonstrating no difference between young and old cell types. Young muscle (SCs) in mono-culture demonstrated greater proliferation at days 7, 10 and 14 of differentiation compared to young muscle paired with either young or old bone (MSCs) cells (p < 0.05, Fig. 2A). This pattern continued in young bone (Fig. 2B), old muscle (Fig. 2C) and old bone (Fig. 2D). Furthermore, no differences were found in proliferation between young and old. Young monocultures were similar to old monocultures in proliferation, as well as co-cultures. Additionally, there were no differences between young bone and old bone, or young muscle and old muscle.

**Figure 2.**
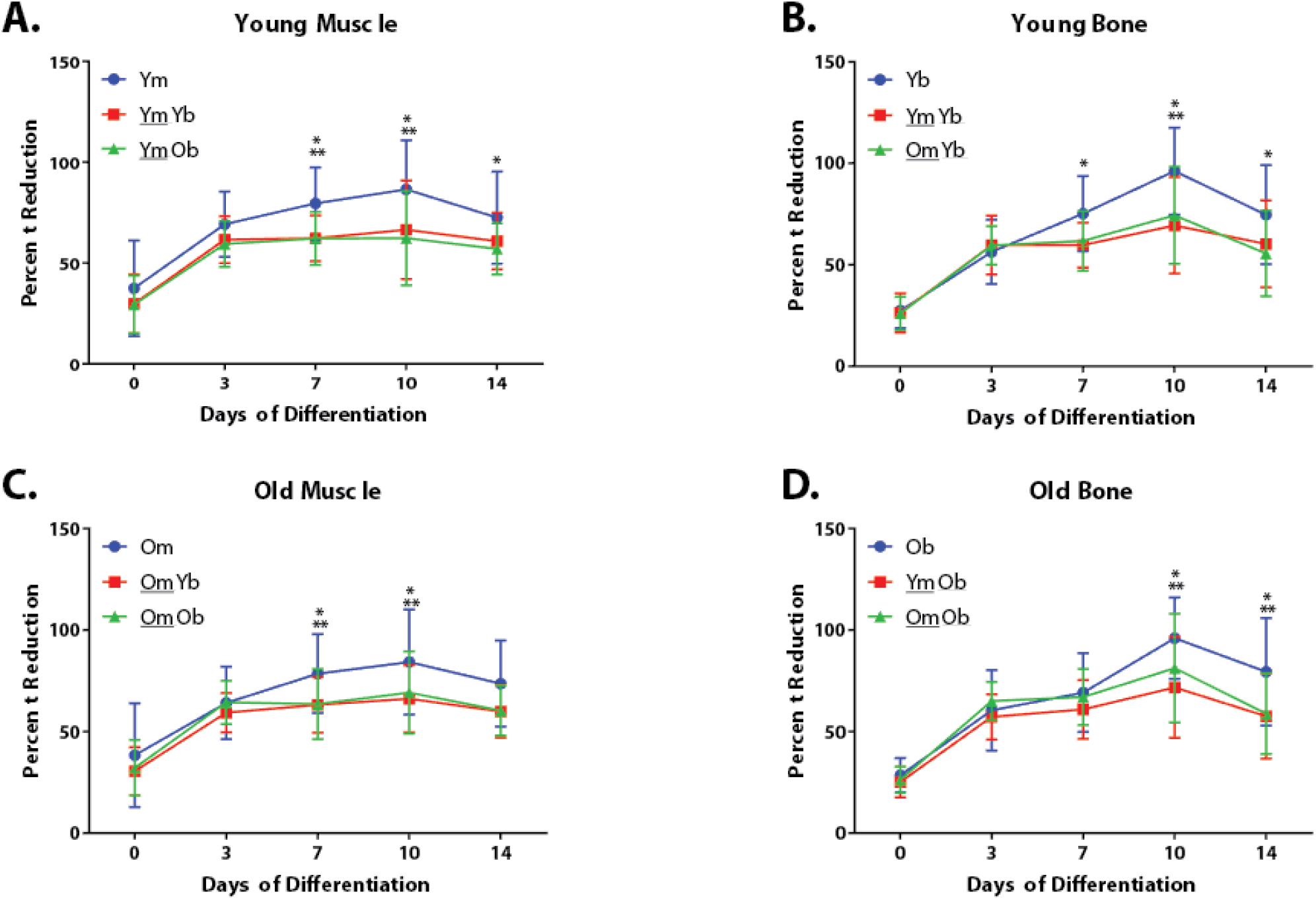
AlamarBlue^®^ reduction across differentiation. Proliferative ability differed between some mono- and co-cultures during differentiation days 7-14 for A) young muscle, B) young bone, C) old muscle, and D) old bone. Data are mean ± SEM. *p < 0.05 for mono-culture (blue) vs. co-culture (red). **p < 0.05 for mono-culture (blue) vs. co-culture (green).

### Collagen and Myosin Content

Collagen immunostaining did not differ across cell or culture type (Fig. 3A). In general, differences in collagen expression between mono-cultures and co-cultures were small (p > 0.05), according to the amount of collagen staining in a given field. When these values were normalized to intensity or number of cells in the area, there still was no difference. For myosin, the relative expression in muscle cells compared to that in bone cells was generally greater in co-culture than in mono-culture (p < 0.05, Fig. 3B), with no difference in the amount of myosin expressed between young and old cells, either in mono- or co-culture (p > 0.05). In mono-culture, myosin expression did not differ between muscle and bone cells regardless of age (p > 0.05). However, myosin expression in muscle cells seemed to trend higher than in bone cells. In co-culture, bone cells generally had little to no expression of myosin relative to their muscle cell counter-parts (p < 0.05). Age had no effect on the amount of myosin expressed in these co-cultures. Similarly, when controlling for intensity or number of cells within the field, these differences remained the same (data not shown). Representative images of these condition are shown in Figures 3 C, D, and E.

**Figure 3.**
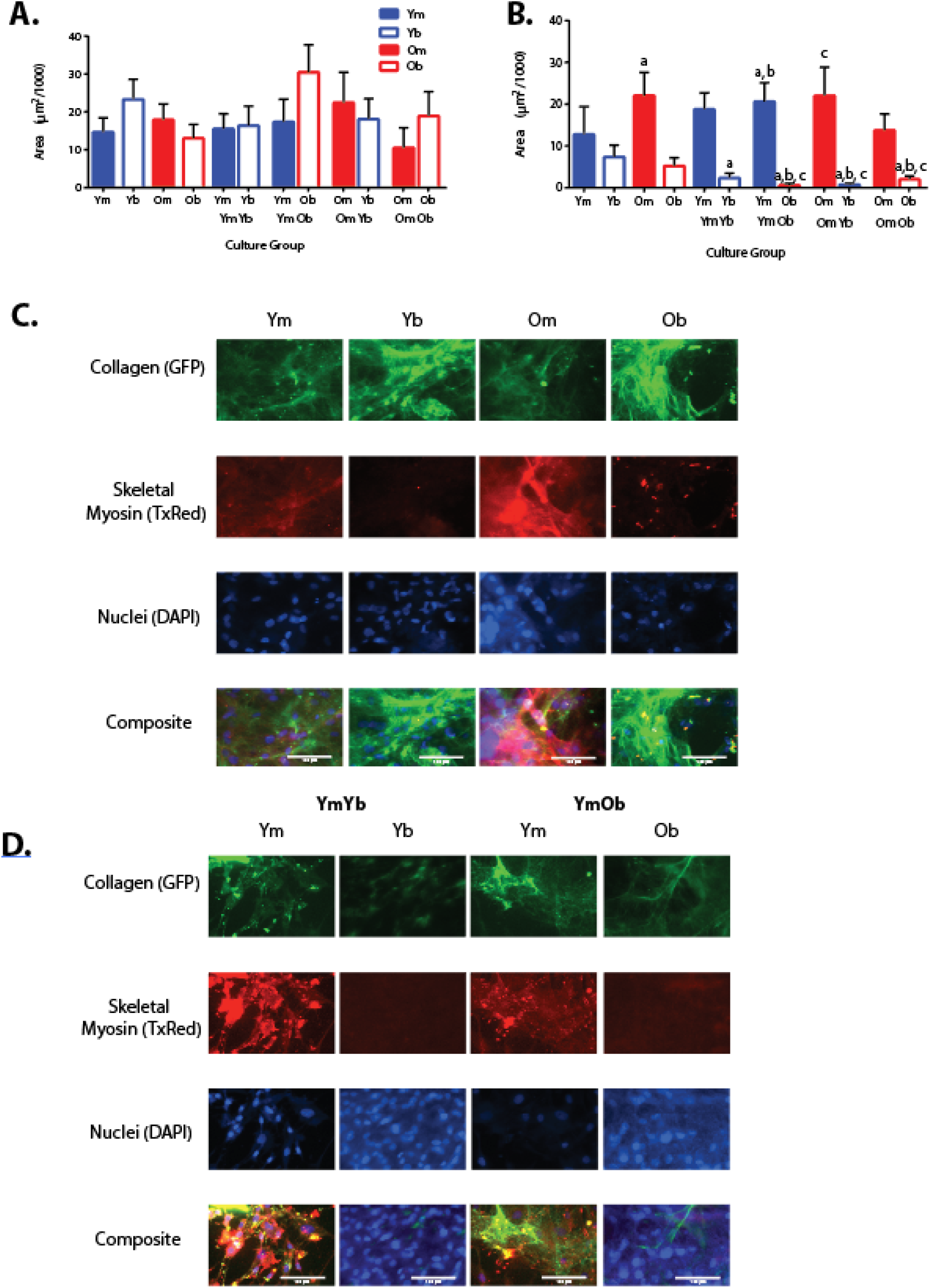
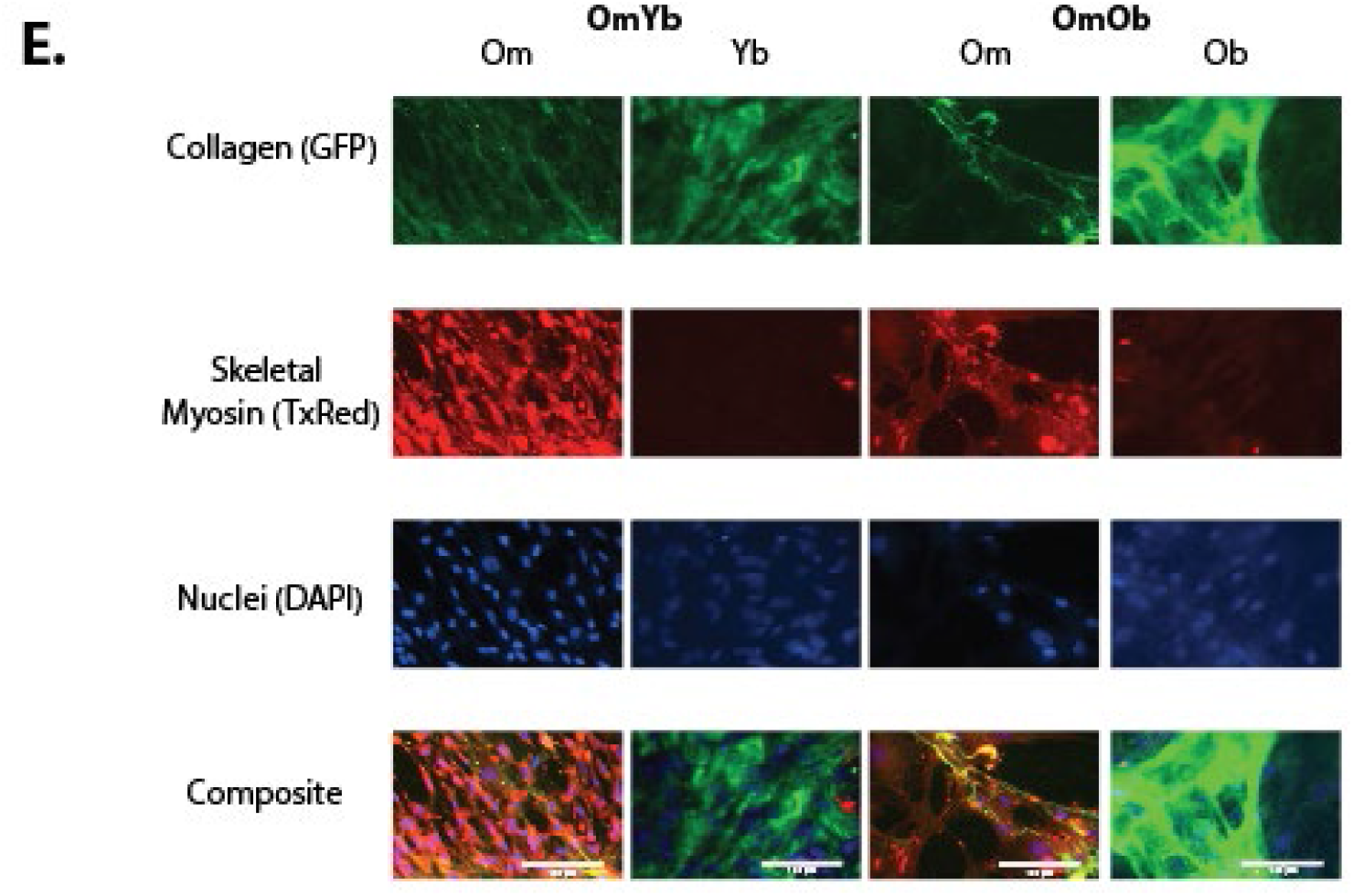
Immunocytochemistry analysis of cultured cells. Collagen I expression (A) and skeletal myosin expression (B) after 14 days of differentiation. No differences in collagen expression were found across cell type or age, or in mono-vs. co-culture (p < 0.05). Myosin expression was typically increased in SCs in co-culture (*,p < 0.05) and tended to be increased compared to MSCs in mono-culture. Age had no effect between these conditions. Representative images from the various conditions are shown in (C), (D), and (E). Data are mean ± SEM.

### Mineralization results

Alizarin red results in mono-culture showed that bone cells had or tended to have significantly more amounts of mineralization relative to muscle cells (p < 0.05), yet age did not have a significant effect (p > 0.05, Fig. 4A). In co-culture, however, all cell types and ages had very little mineralization (p > 0.05). Between mono- and co-cultures, mineralization was higher in bone cells in mono-culture than in co-culture. These data suggests that osteoblasts more successfully differentiated into osteoblasts in mono-culture than in co-culture.

**Figure 4.**
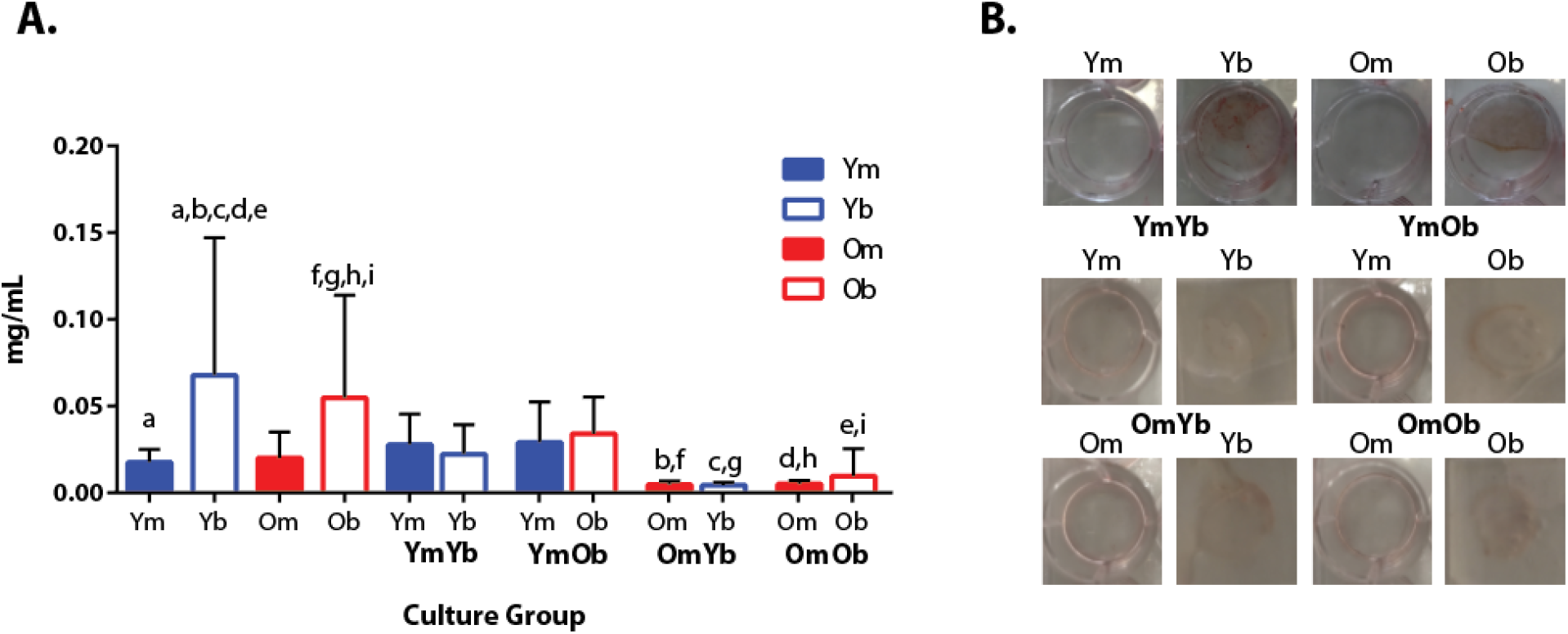
Alizarin red solubilization data from fixed cells. Solubilization results (A) and representative images (B). In mono-culture, MSCs tended to have or had a significantly greater amount of mineralization compared to SCs (p < 0.05). In co-culture, MSCs paired with old SCs had significantly less mineralization compared to their mono-culture equivalent (p < 0.05). MSCs paired with young SCs had similar amounts of mineralization compared to mono-culture (p < 0.05). Data are mean ± SD. Bars with the same letter above them are significantly different.

### IGF-1 Content

ELISA results for IGF-1 expression in conditioned media demonstrated no difference between cell or culture type (Fig. 5). In mono-culture, bone cells tended to have lower expression of IGF-1 compared to muscle cells, but these differences were not significant; young and old cells had similar IGF-1 expression in mono-culture. In co-culture, IGF-1 expression did not differ between cell types or between young and old cells. Furthermore, no differences were found in the amount of expression between mono- and co-cultures. These results suggest that IGF-1 expression is not altered by communication between these two cell types in aging.

**Figure 5.**
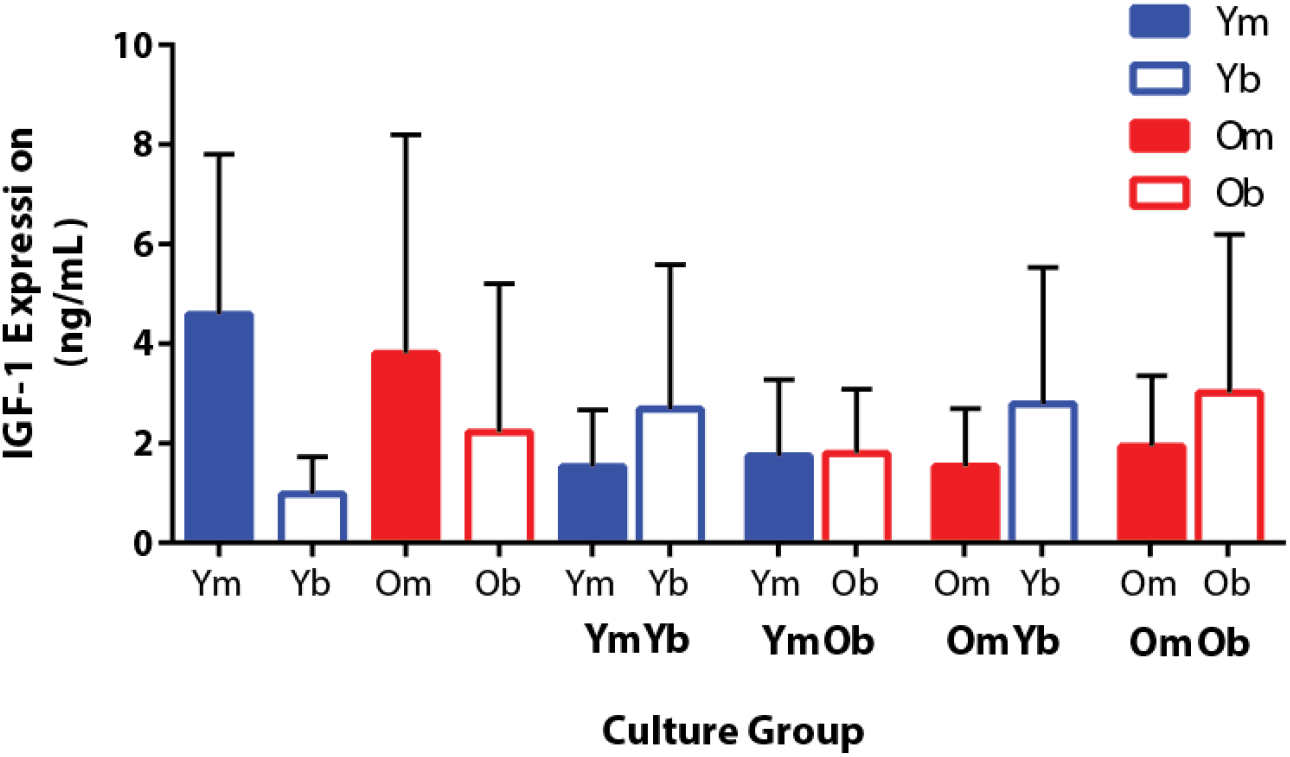
IGF-1 expression from conditioned media on day 14. IGF-1 expression did not differ by cell type, culture type, or age (p > 0.05). Data are mean ± SEM.

## Discussion

Understanding the bone-muscle interactions has become increasingly important to better understand musculoskeletal diseases. Understanding these interactions with regards to aging could lead to novel therapeutic options to improve aging outcomes. Here, we utilized a previously optimized co-culture system to investigate the bone-muscle interactions in both young and old cells. Our hypothesis was that young cells would generally demonstrate greater proliferation and differentiation compared to old cells. Furthermore, when placed in co-culture, young cells paired together would experience increased crosstalk between the cell types; young cells paired with old cells would be able to improve the old cells’ proliferation, differentiation, and crosstalk; and old cells paired together would perform the worst in regard to proliferation, differentiation, and crosstalk. The results of these experiments demonstrated that our hypotheses were partially supported.

### Cellular proliferation

Our primary hypotheses for cellular proliferation predicted that young cells would proliferate more than old cells, and cells in co-culture would synergistically undergo greater proliferation compared to the mono-cultured cells. Neither of these hypotheses were supported by the alamarBlue^®^ results, which showed no difference in proliferation between young and old cells and greater proliferative ability at later days of differentiation for mono-cultures compared to co-cultures. Other studies investigating the proliferative potential between two cell types have typically demonstrated a greater proliferation response for co-cultures compared to mono-cultures.^23,24^ The differences between these finding and our own, however, lie in the selection of animal model, cell type, and type of co-culture. In our case, we are the first to use this animal model of aging in a co-culture study between SCs and MSCs. The decreased proliferation found in our co-cultures may be the result of a regulatory effect between the two cell types in co-culture. This response may mimic those found in the *in vivo* state, where bone-muscle interactions can restrict the growth of both tissue types.^25^ More interestingly, however, is that this restrict effect was conserved across age in our data. Aging can hinder many aspects of the cellular lifecycle, such as a characteristically reduced proliferative capacity.^26-28^ Our old cell data did not demonstrate this reduced proliferation compared to young cells and may indicate a less severe aging phenotype in our cells than present in other studies. This less severe phenotype was observed not only in proliferation but many of the cellular activities in our data set.

### Cellular differentiation

For differentiation, our hypotheses were 1) old SCs would have a lower capacity to differentiate (measured by skeletal myosin deposition) compared to young SCs; 2) SCs overall would have larger amounts of myosin compared to MSCs; 3) co-cultured SCs would synergistically, due to the presence of MSCs, have more myosin deposition compared to mono-cultured SCs; and 4) the presence of young MSCs would induce old SCs to produce greater amounts of myosin compared to old SCs cultured with old MSCs. Here, our hypotheses were only partially supported. In co-culture particularly, SCs had greater myosin content compared to MSCs, yet age had no significant effect. Other studies have shown that the amount of myosin production decreases in old SCs as they become exhausted, less proliferative, and shift to producing more collagen or other fibrotic material instead.^29,30^ In mono-culture, myosin expression was not significantly different between SCs and MSCs, although SC myosin expression tended to be greater. These data suggest that, in co-culture, MSCs may have a beneficial effect on the differentiation of SCs. Another study utilizing co-culture in other cell types (e.g., MSCs and cardiomyocytes) also demonstrated this synergistic response, suggesting that crosstalk signals between two cell types may be the source of enhanced differentiation.^31^ Taken together with the viability results, this suggests that instead of actively dividing, the SCs in co-culture conditions favored differentiation more so than in mono-culture. In the future, a more morphological investigation of myoblast fusion could provide a more definitive explanation for differentiation changes as described by others previously. Morphological images taken at time points throughout differentiation could demonstrate shifts in the timing of myoblast fusion with age.

Surprisingly, MSC differentiation showed an opposite response compared to SCs. MSCs demonstrated greater mineral deposition, indicative of increased differentiation, in mono-cultures compared to co-cultures, with little effect of age. We hypothesized that MSCs would have more mineralization in co-culture, with greater effects in young cells, neither of which were supported by our data. We posit two explanations for these observations: 1) SCs inhibited MSC differentiation or 2) media mixing between the two wells diluted differentiation media in MSC wells or a combination of the two. In the first case, experiments exploring bone-muscle crosstalk using isolated monocultures and conditioned media have demonstrated that signaling factors from muscle, such as myostatin and ciliary neurotrophic factor 1 (CNF-1), inhibit osteoblast differentiation.^32,33^ These factors are secreted by muscle after sufficient growth occurs to inhibit further muscle growth. In our preliminary studies and those of others, SCs have been observed to reach myoblast fusion as early as 10 days,^34-36^ whereas MSCs differentiation can take as long as 21 days in some studies^37,38^. We suggest that SCs were reaching a more differentiated state more quickly than MSCs and may have secreted these inhibitory factors before MSCs could completely differentiate. We also found that growth of myoblasts occurred more quickly than MSCs and would result in monolayers lifting off the plate after 14 days, precluding us from continuing to 21 days to see if MSCs could reach a more differentiated state. In the second case, the Transwell^®^ setup required that each cell type receive its own differentiation media, yet the pores within the Transwell^®^ did not inhibit mixing between the two wells. While another study reported that this mixing can take several days^39^, and may not have been a factor in this study, we cannot exclude this possibility. In either case, the presence of significant mineralization in mono-cultures suggest that that the differences have arisen due to the co-culture setup itself. Furthermore, the lack of difference between young and old cultures continues to reinforce that the cells developed a less severe aging phenotype.

### Cellular crosstalk

To investigate crosstalk, we investigated the role of IGF-1 in signaling between the two cell types, as well as collagen I expression in the two cell types. With age, muscle commonly loses its ability to repair and replace tissue effectively and instead deposits collagen. Additionally, bone experiences collagen loss, resulting in an increased incidence of fractures and microcracks with age.^9^ IGF-1 is a potent growth factor of both muscle and bone and can enhance collagen production. We hypothesized that, with age, IGF-1 expression would be decreased, leading to decreases in collagen I deposition in both SCs and MSCs. In co-culture experiments, we hypothesized that old cells plated with young cells would behave more similarly to young cells and have similar IGF-1 and collagen I expression, but our results did not support these hypotheses. Instead, our results show that IGF-1 expression (Fig. 5) and collagen expression (Fig. 3A) did not differ across the culture conditions, cell types, or age. Many studies have highlighted the role of crosstalk between muscle and bone and the importance of this crosstalk for the growth and development of both.^5^ In this study, we focused on IGF-1, as it is an important mediator of growth and is often reduced with age.^40^ However, IGF-1 has only been minimally investigated during bone-muscle crosstalk.

Our results with no age-related changes in IGF-1 expression were surprising, considering the differences in IGF-1 expression observed in several aging studies, but this discrepancy may result from the differences in study design such as using *in vitro* experiments instead of *in vivo* experiments. IGF-1 expression is frequently measured in serum samples^41^ or in whole tissue samples.^42^ While both bone and muscle can secrete and react to their own IGF-1 signals in an autocrine fashion, perhaps the IGF-1 expression in bone and muscle in an *in vivo* environment are enhanced by additional IGF-1 from the bloodstream and elsewhere. In the isolated *in vitro* environment used here, we found that IGF-1 is clearly secreted by bone and muscle but may only promote initial cell growth and not the continued growth and differentiation. Therefore, while we did not observe differences in IGF-1 expression associated with age or co-culture conditions in this isolated system, it cannot capture the system-wide IGF-1 signaling that occurs *in vivo*. Furthermore, IGF-1 is merely one bone-muscle crosstalk factor that can be detrimentally affected by aging; studies of additional signaling factors could provide a more complete picture of age-related crosstalk changes *in vitro*.^43-45^

IGF-1 was also investigated due to its significant role in downstream collagen production. We anticipated diminished amounts of IGF-1 expression with aging would lead to decreased collagen deposition in old cells compared with young cells. In this context, the lack of differences in collagen I expression between cell types, ages, or culture types may not be so surprising. In typical *in vivo* environments, bone has approximately 36% of its composition as collagen I, and this content decreases with age.^46,47^ Conversely, muscle has 2-6% collagen I, and content increases with age.^48,49^ However, collagen I deposition *in vitro* was not different between bone and muscle cells. Since collagen is deposited by mature cells, and given the early maturation stage of our cells, perhaps the differentiation time was not sufficient to allow significant collagen deposition. Furthermore, in co-culture, the lack of difference in collagen deposition may be indicative of a lack of influence of the crosstalk between the two cell types. While differences in IGF-1 expression or collagen deposition were not observed with co-culture, we cannot exclude the possibility that other signaling pathways or protein expression were not altered by crosstalk. Further studies are required to probe this possibility.

## Conclusions

In summary, we demonstrated that co-culture of bone and muscle cells regulates proliferation more so than for cells grown in mono-culture, yet age or cell type did not affect proliferation. Furthermore, differences in cellular differentiation or crosstalk were minimal, with little to no effect of age, cell type, or culture condition on myosin expression, mineralization, IGF-1 expression, or collagen deposition. This study was the first to use the F344 x BN F1 hybrid rat in a study for cellular aging while also investigating the crosstalk between SCs and MSCs. The lack of differences between young and old cellular proliferation, differentiation, and crosstalk demonstrate that this particular model may not be ideal for age-related cellular studies. However, the use of this model to successfully grow SCs and MSCs should provide evidence to continue using this system for *in vitro* aging crosstalk studies. The use of a Transwell^®^ co-culture system provides a framework with which to understand the individual contributions of each cell type, which cannot be isolated in direct co-culture models. Many potential bone-muscle crosstalk factors besides IGF-1 could be contributing to the concomitant age-related changes observed in bone and muscle *in vivo*, and future studies should focus either on developing *in vitro* systems that better mimic the *in vivo* aging environment and/or examining contributions of different signaling pathways to tissue aging. Overall, this study demonstrated that the intricate interactions between bone and muscle depends on a variety of cues, and advancing our understanding of these cues is essential for developing better models for mimicking aging and better treatments to mitigate functional musculoskeletal deficits with aging.

